# Communication between the nucleus and the mitochondria via NDUFS4 alternative splicing in gastric cancer cells

**DOI:** 10.1101/2022.10.07.511259

**Authors:** V. Papadaki, Z. Erpapazoglou, M. Kokkori, M. Rogalska, E. Tsakiri, H. Akhtorab, D. T. Smoot, K. Papanikolopoulou, M. Samiotaki, P. Kafasla

**Author notes:** equal contribution (V. Papadaki, Z. Erpapazoglou, M. Kokkori).

## Abstract

A constant communication between the nucleus and the mitochondria allows both organelles to ensure cellular homeostasis and adaptation to mitochondrial stress. Mitochondrial biogenesis and function are controlled by anterograde regulatory pathways involving a large number of nuclear-encoded proteins. Transcriptional networks controlling the nuclear-encoded mitochondrial genes are known, however alternative splicing (AS) regulation has not been implicated in this communication. Here, we show that IQGAP1, a scaffold protein that regulates AS of distinct subsets of genes in gastric cancer cells, participates in AS regulation that strongly affects mitochondrial respiration. Combined proteomic analyses and RNA-seq profiles of *IQGAP1*^*KO*^ and parental cells show that *IQGAP1*^KO^ alters a specific AS event of the mitochondrial respiratory chain complex I subunit NDUFS4 and downregulates a subset of complex I subunits. In *IQGAP1*^KO^ cells, respiratory complex I intermediates accumulate resembling assembly deficiencies observed in patients with Leigh syndrome bearing *NDUFS4* mutations. Mitochondrial complex I activity is significantly lower in KO compared to parental cells, while exogenous expression of IQGAP1 partially restores *NDUFS4* AS pattern and expression and reverses mitochondrial defects of *IQGAP1*^KO^ cells. Our work sheds light to a novel facet of IQGAP1 in mitochondrial quality control that involves fine-tuning of complex I activity through AS regulation.

## Introduction

Cancer cells use multiple stress response pathways to counteract exogenous or endogenous stressors. Such a stress response involves anterograde (nucleus to mitochondria) and retrograde (mitochondria to nucleus) signalling between mitochondria and the nucleus in an attempt to control mitochondrial function. Specifically, mitochondrial function is controlled by the nucleus through anterograde signals, which allow these organelles to adapt to the cellular environment^1^. This regulation is known to be based mainly on the expression of nuclear-encoded mitochondrial proteins that induce mtDNA gene expression, and on several transcription factors and co-regulators that regulate the expression of the nuclear-encoded mitochondrial proteome. Nothing is known to date for a role of AS in anterograde signalling.

In humans, the importance of accurate AS in health and disease, including cancer, has been well documented^2–4^. Cancer cells have general as well as cancer type-specific and subtype-specific alterations in the splicing process that can have prognostic value and contribute to every hallmark of cancer progression, including deregulation of cellular energetics^5^. Compared with normal cells, tumour cells preferentially metabolize glucose through aerobic glycolysis, and some of the genes encoding enzymes involved in aerobic glycolysis are prone to alternative splicing. A number of studies have unveiled some connections between glucose metabolism and splicing programs. A well-studied example is pyruvate kinase: alternative splicing modulates the ratio between M1 and M2 isoforms of PKM and thus determines the choice between aerobic glycolysis and complete glucose oxidation in the Krebs cycle^6^. Reciprocally, intermediates in the Krebs cycle may impact splicing programs at different levels by modulating the activity of 2-oxoglutarate-dependent oxidases^7^. AS has been also shown to affect glutamine and fructose metabolism by altering the expression of splicing variants of enzymes involved in these processes^8^. However, an important role of respiration—and mitochondrial metabolism more broadly—in cancer cell and tumour growth is becoming more and more apparent and several inhibitors of oxidative phosphorylation and related pathways have now advanced into clinical trials^9^. Even though a connection between AS and glucose metabolism has been established, the link between AS and mitochondria respiration is just emerging as a crucial investigation-demanding field^10^.

Recently, we identified a tumour promoting AS program, supported by a nuclear pool of the scaffold protein IQGAP1 (IQ Motif Containing GTPase Activating Protein 1) and its interacting partners that are components of the spliceosome machinery^11^. We showed that in gastric cancer cells IQGAP1 participates in the regulation of AS of a number of cell cycle related pre-mRNAs, thus promoting tumour development. IQGAP1 does this by affecting the post-translational modification level and/or subnuclear distribution of certain splicing factors.

Given that nuclear IQGAP1 is differently distributed in the nuclei of gastric cancer cells of different origin^11^, we studied further the consecutive different AS regulation by IQGAP1. We present herein conclusive evidence on the participation of IQGAP1 in the oxidative metabolism of gastric cancer cells via its involvement in the regulation of AS of components of the mitochondrial electron transport chain.

## Materials and Methods

### Reagents

Unless stated otherwise, all chemicals were purchased from Sigma-Aldrich or ThermoFisher Scientific. The following antibodies were used: anti-β-actin (Novus Biologicals, Cat# NB600-501), anti-ATP5A (Abcam, clone 15H4C4, Cat# ab14748), anti-COX4 (Novus Biologicals, Cat# NB110-39115), anti GAPDH (Protein Tech, Cat# 60004-1-Ig), anti-HSP60 (Santa Cruz, clone B-9, Cat# sc-271215 or Protein Tech, Cat# 15282-1-AP), anti-IQGAP1 (Santa Cruz, clone D-3, Cat# sc-374307 or clone C-9, Cat# sc-376021), anti-Lamin B1 (Santa Cruz, clone A-11, Cat# sc-377000), anti-Myc-Tag (Cell Signaling, clone 9B11, Cat#2276), anti-NDUFS4 (Novus Biologicals, Cat# NBP1-89026-25), anti-NDUFA4 (Santa Cruz, clone 2G7, Cat# sc-517091), anti-NDUFS1 (Santa Cruz, clone E-8, Cat# sc-271510), anti-NDUFAF2 (mimitin) (Santa Cruz, clone H-11, Cat# sc-365592), anti-SDHA (Santa Cruz, clone F-2, Cat# sc-390381), anti-UQCRC2 (Santa Cruz, clone G-10, Cat# sc-390378), anti-VDAC2 (Protein Tech, Cat# 11663-I-AP), HRP-conjugated goat anti-rabbit (SouthernBiotech, Cat# 4050-05), HRP-conjugated goat anti-mouse IgG (SouthernBiotech, Cat# 1030-05), anti-rabbit-Alexa Fluor 555 (Molecular Probes, Cat# A27039), anti-rabbit-Alexa Fluor 594 (Molecular Probes, Cat# A21207), anti-mouse Alexa Fluor 488 (Molecular Probes Cat# A28175).

### Cell cultures

Human gastric cancer cell lines MKN45 and NUGC4 and the non-cancerous gastric epithelial cell line HFE145 were used for this study. MKN45 is a human gastric adenocarcinoma cell line derived from liver metastatic tissue, while NUGC4 originates from a proximal metastasis in paragastric lymph nodes. MKN45 and NUGC4 cells were cultured in RPMI medium (GIBCO Cat# 31870-025) and HFE145 cells in DMEM medium (Lonza, Cat#12-733F), supplemented with 10% FBS, 1% L-glutamine and 1% penicillin-streptomycin, under standard tissue culture conditions (37^0C^, 5% CO2). *IQGAP1*^KO^ cells for all the above cell lines were generated with the CRISPR/Cas9 approach, as previously described^11^.

NUGC4 parental and KO cells were transfected with 4-8 μg of pcDNA3.1 or pcDNA3.1-myc-IQGAP1, using Turbofect (Thermo Fisher Scientific, Inc., MA) or FuGENE® 6 (Promega Cat# E2961) as transfection reagents, according to the manufacturer’s protocol. Alternatively, the cells were transfected with 10 μg plasmid by electroporation, using the Super Electroporator NEPA21 (Nepa Gene Co., Ltd; Poring pulse: pulse voltage 175 V; pulse interval 50 ms; pulse width 5 ms; pulse number 2; and Transfer pulse: pulse voltage 20V; pulse interval 50 ms; pulse width 50 ms; pulse number 5). Note that under all tested conditions the transfection efficiency estimated by the % of myc-IQGAP1-expressing cells was around 10%. Subsequent analyses were performed 48 h after transfection.

### RNA isolation, cDNA synthesis and PCR/qPCR

Total RNA was extracted with the TRIzol® reagent (Thermo Fisher Scientific, Inc). Residual contaminating DNA was removed with DNase I (New England BioLabs, Inc.) treatment. Reverse transcription was carried with 1μg of total RNA, using random hexamer primers (Invitrogen), oligodT18 (New England BioLabs, Inc.), RNaseOUTTM Recombinant Ribonuclease Inhibitor (Thermo Fisher Scientific) and Protoscript II reverse transcriptase (New England BioLabs, Inc.), according to manufacturer’s instructions. qPCR was performed in a Corbett Rotor -Gene™ RG 6000 using Kapa SYBR®FAST (KAPA Biosystems) qPCR mix. The PCR samples were loaded onto 8% acrylamide-urea gel. Quantification of % exon inclusion was performed using ImageJ.

### RNA-seq analysis

cDNA libraries for NUGC4 and NUGC4-*IQGAP1*^KO^ cells were prepared in collaboration with Genecore, at EMBL, Heidelberg. Alternative splicing was analyzed by using VAST-TOOLS v2.2.2 and expressed as changes in percent-spliced-in values (ΔPSI). A minimum read coverage of 10 junctions reads per sample was required. Psi values for single replicates were quantified for all types of alternative events. Events showing splicing change |ΔPSI | > 15 with minimum range of 5% between NUGC4 and KO samples were considered IQGAP1-regulated events.

### Western blot

15-30 μg of protein extract were resolved on an 8% or 12% polyacrylamide-SDS gel and transferred to a polyvinylidene difluoride membrane (PVDF, Millipore). Primary antibodies were used at the recommended dilutions in 5% milk or 3% BSA diluted in TBS-Tween. As secondary antibodies were used an HRP-conjugated goat anti-mouse IgG or an HRP-conjugated goat anti-rabbit IgG (1:5000). Detection was carried out with the Immobilon Crescendo Western HRP substrate (Merck Millipore®), using the Chemidoc imaging system and the Image Lab software (Bio-Rad Laboratories).

### Immunofluorescence and mitochondrial network analysis

Cells were next for 10 minutes with 4% paraformaldehyde in 1X PBS (phosphate buffer saline), followed by permeabilization with 1X PBS-0.25% (v/v) Triton X-100. After blocking for 45 min with 5% BSA in 1X PBS/0.25% Triton X-100/, cells were incubated with primary antibodies mouse anti-Myc (1:2000) and/or rabbit anti-HSP60 (1:3000). The secondary antibodies used were anti-mouse-Alexa Fluor 488 and anti-rabbit Alexa Fluor 555 or 594 (1:500). Nuclei were stained with DAPI (1 μg/ml). Images were acquired on a LEICA SP8 White Light Laser confocal microscope and were analyzed using Image J. Mitochondrial network morphology was analyzed using the Mitochondria Analyzer plugin (Ahsen Chaudhry doi: 10.1152/ajpendo.00457.2019), using the following settings: 2D threshold, mean with block size at 1.450 microns and C-value at 5.

### Mitochondrial respiratory chain complex I activity

Mitochondrial Complex I activity was measured with Complex I Enzyme Activity Microplate Assay Kit (Cat# ab109721, Abcam), according to the manufacturer’s instructions, using 400 μg of total or 35 μg of mitochondria-enriched protein extract. Measurement at 450 nm was carried out on a Sunrise™ (Tecan) instrument over a period of 1800 sec, with a 20 sec interval.

### Mitochondrial ROS levels

Cells were incubated for 30 minutes at 37°C with 0.5 μM MitoSOX Red (Thermo Fisher Scientific, Inc.). They were then collected by trypsinization, resuspended in 1X PBS and analyzed on a FACSCanto™II (BD Biosciences) instrument, before and after addition of Propidium Iodide. The appropriate negative (unstained) and positive (cells treated with 50 μM antimycin A prior to the staining) controls were included. Analysis of MitoSOX Red fluorescence was performed with FlowJo™.

### Mitochondrial mass

Cells were incubated for 20 minutes at 37°C with 0.1 μM MitoTracker Green FM (Thermo Fisher Scientific, Inc.). They were then collected by trypsinization, resuspended in 1X PBS and analyzed on a FACSCanto™II (BD Biosciences) instrument, before and after addition of Propidium Iodide. The appropriate negative (unstained) control was included. Analysis of MitoTracker Green FM fluorescence was performed with FlowJo™.

### Isolation of nuclear and mitochondria-enriched fractions

Nuclear isolation was performed as described^12^. Briefly, cells were re-suspended in 3 to 5 volumes of hypotonic Buffer A (10 mM Tris-HCl, pH 7.4, 100 mM NaCl, 2.5 mM MgCl2) supplemented with 0.5 % Triton X-100, protease and phosphatase inhibitors (1 mM NaF, 1 mM Na3VO4) and incubated on ice for 10 min. Cell membranes were sheared by passing the suspension 4-6 times through a 26-gauge syringe. After centrifugation at 3000 x g for 10 min at 4°C, the supernatant was kept as the cytoplasmic extract, and the nuclear pellet was collected, washed once and resuspended in 2 volumes of Buffer A. The nuclei were sonicated twice for 5s (0.2A). Then, samples were centrifuged at 4000 x g for 10 min at 4^0C^, to separate the nuclear extract (supernatant) from the nuclear pellet. The latter was re-suspended in 2 volumes of 8 M Urea.

Crude mitochondrial fractions were prepared as previously described^13^. Cells were resuspended in mitochondrial isolation buffer (10 mM Tris-MOPS pH 7.4, 200 mM sucrose, 1 mM EGTA-Tris pH 7.4) supplemented with protease and phosphatase inhibitors and were homogenized with a Teflon Dounce. Unbroken cells and nuclei were eliminated by centrifugation at 600 x g for 10 min at 4^0C^, and the supernatant was centrifuged at 7000 x g for 10 min. The mitochondrial pellet was washed once and resuspended in isolation buffer for further analysis of mitochondrial protein levels, complex I assembly and functional assays (respirometry, complex I activity).

Protein concentration in the extracts was assessed using the Bradford assay^14^.

### Blue Native Gel Electrophoresis (BNGE)

For BNGE analysis, mitochondrial pellets were solubilized with digitonin (8 g/g of protein) or DDM (n-Dodecyl-B-D-Maltoside, 1.6 g/g of protein) for 5 min at 4°C. After centrifugation, loading buffer (50mM NaCl, 50mM Imidazole, 1mM EDTA, 10% w/v glycerol, 5% Coomassie G250) was added to the supernatants. The samples were loaded on 3-12% ready-made acrylamide gradient gels (Invitrogen) and separated in the presence of anode buffer (1M Imidazole) and cathode buffer (50 mM Tricine, 7.5 mM Imidazole, 0.02% Coomassie G 250)9. Gels were transferred onto a PVDF membrane and were used for the analysis of mitochondrial respiratory complexes assembly by Western blot.

### Proteomics analysis

The digested samples were subjected to nLC-nESI MS/MS analysis in an Ultimate3000 RSLC chromatographic system coupled to the HF-X Hybrid Quadrupole-Orbitrap mass spectrometer (Thermo Scientific). Analysis was carried out in a data-dependent positive ion mode (DDA). The spray voltage was set to 2.2 kV with a capillary temperature of 270°C. Full scan MS were obtained on the Orbitrap analyzer at a 60,000 resolution with an Automatic Gain Control (AGC) set to 3e6 and maximum injection time 45 ms. Data were obtained in technical duplicates using the Xcalibur software. The mass spectrometer was operated using the lockmass function on.

The raw files were searched and the identified proteins were quantified using Label Free Quantitation (LFQ) in MaxQuant (version 1.6.14.0), using tryptic/P search against the human uniprot protein database (downloaded 19/09/2019). Search parameters included a molecular weight ranging from 350 to 5,000 Da, a precursor mass tolerance of 20 ppm, an MS/MS fragment tolerance of 0.5 Da, and methionine oxidation, deamidation of asparagine and glutamine and protein N-terminal acetylation were set as variable modifications. The protein and peptide false discovery rate (FDR) was set to 1%. The match-between-run and second peptides functions were enabled. Statistical analyses and visualization (heatmap) were performed in the environment of Perseus software platform.

### Wound healing assay

Cells were cultured in 24-well plates until nearly 90% confluency. Then, scratches were made with a sterile 200 μl pipette tip and fresh medium without FBS was added. The migration of cells in the same wound area was visualized at 0, 24, 48 and 72 hours on an inverted microscope.

### Gene Ontology

Enrichment for GO terms was analysed using ShinyGO v0.76.1 with P value cut-off (FDR) set at 0.05.

### Respirometry analysis of intact cells and isolated mitochondria

Real-time cell metabolic profiling of gastric cell lines was performed with a Seahorse XF HS Mini Analyzer (Agilent). Cells were seeded at the following densities: HFE145 at 104 cells/well, MKN45 at 1.6*104 cells/well and NUGC4 at 1.8*10^4^ cells/well. The next day metabolic analysis was performed using the Agilent Seahorse XF Cell Mito Stress Test Kit with the only modification that cells were initially equilibrated in glucose-free medium and glucose was added at the first step of the analysis, to monitor basal glycolysis. Then oligomycin (1 μM), an inhibitor of mitochondrial ATP synthase, was added to measure oxygen consumption that is linked to ATP production, followed by addition of the uncoupler FCCP (2 μM), to achieve maximal respiration rate, and finally the complex I/III inhibitors rotenone/antimycin A (0.5 μM) to abolish oxygen consumption due to mitochondrial activity. After the analysis, cells were fixed and stained with 0.5% crystal violet, the dye was extracted from the cells, and Oxygen Consumption Rate (OCR) and (extracellular acidification rate) ECAR values were normalized against its absorbance at 595 nm. Respiratory and glycolytic activities (Fig. S1C,E) were calculated as proposed by the manufacturer and described in Yepez et al^15^.

Respiration of isolated mitochondria was determined using a Clark-type oxygen electrode connected to a computer-operated Oxygraph control unit (Hansatech Instruments, Norfolk, UK), as previously described^16^. In brief, 500 μg freshly isolated mitochondria were added to the respiration buffer (120 mM KCl, 5 mM KH2PO4, 3 mM Hepes, 1 mM EGTA, 1 mM MgCl2 and 0.2% BSA, pH 7.2) containing 5 mM glutamate/2.5 mM malate. Basal O2 consumption was recorded (State 2) and 500 μM of ADP were added (State 3; indicates rate of ATP production), followed by the addition of 6 μM of the ATP synthase inhibitor oligomycin (State 4; denotes coupling) and 100 nM of the uncoupler carbonyl cyanide p-trifluoromethoxyphenylhydrazone (FCCP) (State FCCP, maximal respiration). For all the experiments, the temperature was maintained at 25°C and the total reaction volume was 300 μl.

## Results

### IQGAP1 has a discrete regulatory role in AS and in the metabolic profile of gastric cancer cell lines of different types

Recently, we reported the participation of the scaffold protein IQGAP1 in AS regulation in gastric cancer cells through interaction with spliceosomal components and mostly hnRNPM (heterogeneous nuclear ribonucleoprotein M). We showed that in both MKN45 and NUGC4 gastric cancer cell lines, there is a nuclear pool of IQGAP1 that interacts with spliceosomal components^11^. However, the amount and the subnuclear distribution of IQGAP1 is different between NUGC4 and MKN45 cells (***Figure S1A***). Furthermore, the interaction with hnRNPM, and possibly with other splicing factors, is partly RNA dependent in NUGC4 but RNA-independent in MKN45^11^. Given these findings and to address the possibly different role of nuclear IQGAP1 in the two cell lines, we performed RNAseq and analysis for AS differences between NUGC4 and NUGC4-*IQGAP1*^*KO*^ cells^11^ (**Table S1**). We detected 168 differentially spliced events in the absence of IQGAP1 with ∼50% of them being alternative exon events (**Table S1**). Interestingly, the majority of these exons were down-regulated in the absence of IQGAP1 (**Figure S1B**). When we compared these 168 AS events to the AS events that are different between MKN45 and MKN45-*IQGAP1*^*KO* 11^ and even though *IQGAP1* deletion affected a similar number of AS events in both NUGC4 and MKN45 cell lines, there were no events changing in both cell lines. These results indicate a possibly different role for nuclear IQGAP1 in alternative splicing regulation in the two gastric cancer cell lines.

The finding that IQGAP1-dependent AS is different in the two cell lines possibly stems from the fact that the two cell lines are of a different origin: NUGC4 is a cell line derived from a paragastric lymph node metastasis of a poorly differentiated intestinal adenocarcinoma of the signet-ring cell type and has been recently classified as a cell line of the Genomically Stable (GS) subtype of gastric adenocarcinomas; MKN45 is a cell line derived from liver metastasis of a diffused, poorly differentiated adenocarcinoma and has been classified as a cell line of the CIN (Chromosomal Instability) subtype^17,18^. To better understand the difference between the two cell lines, we compared them at the level of total proteome. The proteins that were detected at significantly different levels between the two cell lines clustered in two large groups (**Figure 1A, left panel, Table S2)**. The proteins that were enriched in MKN45 cells mostly correspond to genes participating in the assembly of RNP (mainly ribosomal) complexes and mRNA metabolic processes (mainly translation of mRNA; **Figure 1A, right panel and Table S3**). The proteins that were enriched in NUGC4 cells correspond to genes participating in mitochondria/mitochondrial matrix and the process of generation of precursor metabolites and energy and mostly aerobic respiration (**Figure 1A, right panel and Table S3)**.

**Figure 1.**
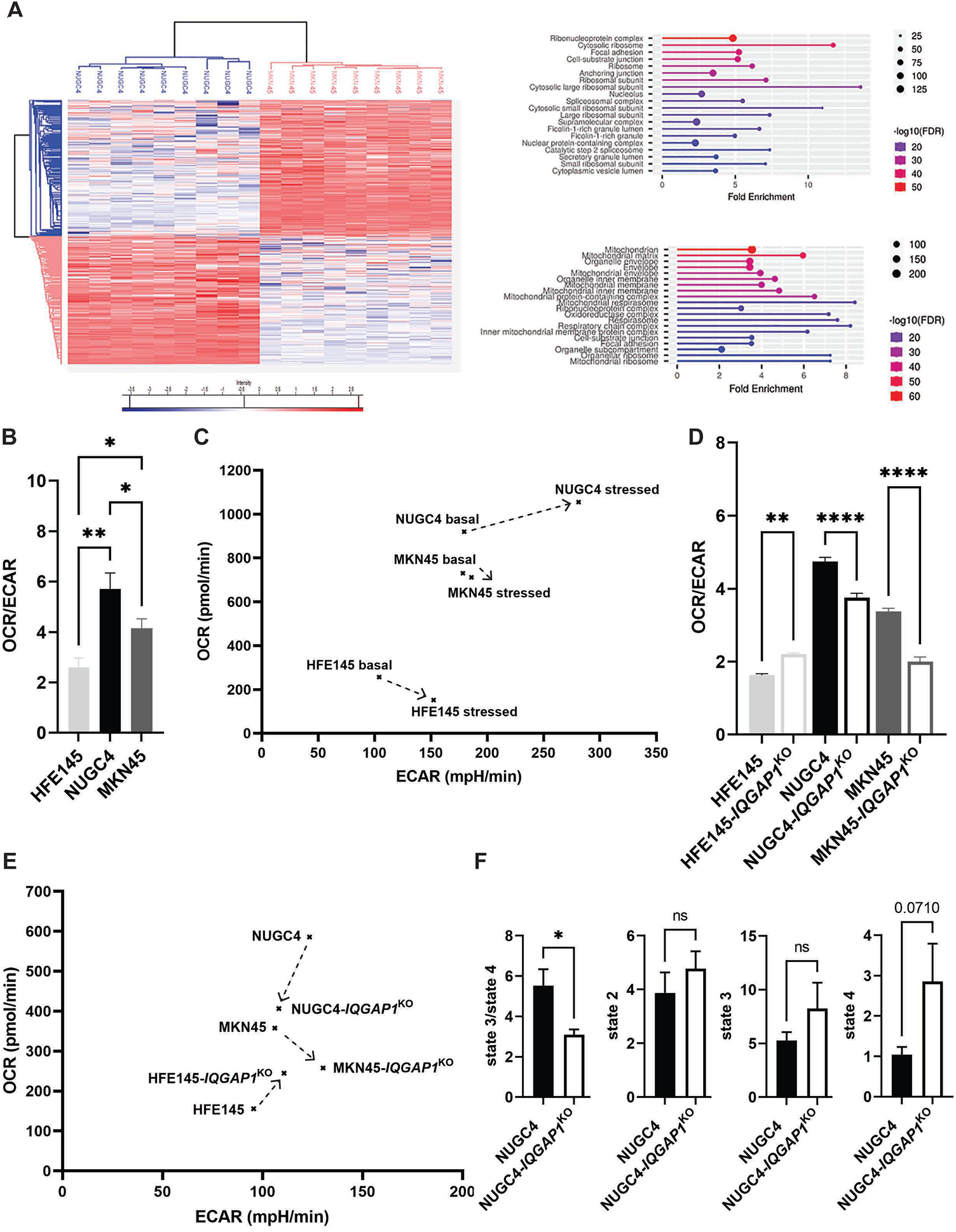
Differential contribution of IQGAP1 to the metabolic profile of specific gastric cancer and normal cell lines. A. Left: Heat map of significantly differentially expressed proteins in NUGC4 and MKN45 cells. Biological triplicates per cell line, subdivided in technical replicates were used for the analysis. Right: Cellular Component Ontology-enrichment analysis of proteins identified as up-regulated in MKN45 (upper panel) or NUGC4 (lower panel) cells. B. Basal OCR to ECAR ratios of HFE145, NUGC4 and MKN45 cell lines measured on a Seahorse analyzer. The mean of 3 independent experiments is presented. C. OCR and ECAR values of the above cell lines plotted at basal (high glucose medium) and stressed conditions (after the addition of FCCP). The arrow indicates the shift in the metabolic profile induced by mitochondrial stress, and reflects the metabolic potential of each cell line, i.e. its capacity to meet an energy demand by increasing oxidative phosphorylation and/or glycolysis. D. Basal OCR to ECAR ratios of *IQGAP1*^KO^ gastric cell lines compared to the parental ones. The mean measurement of 3 independent experiments from each cell line is shown. E. Basal OCR and ECAR values of the KO and parental cells were plotted. Arrows indicate the change in the metabolic potential induced by *IQGAP1*^KO^ for each cell line. F. Respiratory Control Ratio (state3/state4) and absolute respiration rates state 2, state 3 and state 4 of fresh mitochondria preparations from NUGC4 and NUGC4-*IQGAP1*^KO^ cells were measured with a Clark-type electrode. Values represent the mean of 3 independent experiments. Error bars represent ±SEM. *, p<0.05; **, p<0.01; ****, p<0.0001; ns, not significant.

The latter findings were further explored using Seahorse respirometry analysis to compare the metabolic profiles of the two gastric cancer cell lines. NUGC4 and MKN45 cells display an energetic phenotype and rely more on oxidative phosphorylation than glycolysis for energy production (OCR/ECAR of 5.7 and 4.1, respectively), compared to a non-cancerous gastric epithelial cell line HFE145^19^, which was used as a control and was found to rely almost equally on the two metabolic processes (OCR/ECAR of 2.6, **Figure 1B-C, S1C**). Both basal and maximal mitochondrial respiratory capacities of NUGC4 cells are significantly higher than those of MKN45. Moreover, NUGC4 cells show higher metabolic potential than MKN45, reflected by the shift in their metabolic profile under stress, and rely on mitochondrial reserves, as opposed to MKN45 that compensate for energy production by increasing their glycolytic activity (**Figure 1C**). Interestingly, when we checked the contribution of IQGAP1 on the metabolic profile of gastric cancer cells, we found that *IQGAP1*^KO^ impairs their metabolic potential (**Figure 1D-E** and **Figure S1D-E**). Mitochondrial respiratory capacities are significantly lower in both gastric cancer KO cell lines compared to the parental ones, but MKN45 KO cells maintain intact or even increase their glycolytic activity. Interestingly, we observed no major effect of *IQGAP1*^KO^ on the metabolic profile of the non-cancerous HFE145 cells. The effect of IQGAP1 deletion on the respiratory capacity of the NUGC4 cells, which rely the most on mitochondrial respiration was also assessed on crude mitochondria fractions from NUGC4 and NUGC4-*IQGAP1*^KO^ cells using a Clark-type electrode (**Figure 1F**). The Respiratory Control Ratio (RCR, state 3/ state 4) of the KO cells shows a significant reduction (45%) compared to that of the parental cells, resulting from a marked increase in state 4 respiration, which itself is suggestive of an uncoupling of substrate oxidation from ATP production in these cells^20^

Overall, MKN45 and NUGC4 cells display distinct metabolic phenotypes. Though IQGAP1 depletion impairs the metabolic activity of both cancer cell lines, it seems to have a stronger impact on gastric cancer cells that rely highly on mitochondrial respiration affecting possibly the electron transport chain (ETC).

### IQGAP1 affects ETC complex I function via regulation of NDUFS4 levels

To further delineate the effect of IQGAP1 depletion on mitochondria respiration, we analyzed the proteome of crude mitochondrial preparations in *IQGAP1*^KO^ cells and the parental NUGC4 (**Figure S2A**) and MKN45 cell lines. In agreement with our previous observations, IQGAP1 depletion affected differently the mitochondria proteome in the two cell lines: 155 proteins were found to be significantly altered in MKN45-*IQGAP1*^KO^ cells *versus* MKN45, whereas 378 proteins were changed in NUGC4-*IQGAP1*^KO^ (**Table S4**). GO term enrichment analysis of the differentially expressed proteins upon IQGAP1 depletion in NUGC4 cells revealed that they are enriched in proteins involved in the biological process of mitochondrion organization (FDR: 9.98E-11) and in particular mitochondrial respiratory chain complex I assembly (FDR: 1.47E-10) (**Table S4**). More specifically, a large number of protein components of the mitochondrial respiratory complex I are downregulated in NUGC4-*IQGAP1*^KO^ cells compared to the parental ones (**Figure S2B**), but these and other components of complex I are unaffected by the depletion of IQGAP1 in MKN45 cells (**Table S4**). Instead, in MKN45 cells, IQGAP1 deletion results mainly in differentially expressed proteins largely involved in the biological process of actin cytoskeleton organization (FDR: 9.23E-8).

To gain insight into the molecular mechanism by which IQGAP1 regulates mitochondrial respiration, and to address the possible role of AS in such function, we compared the list of significantly changed genes in NUGC4 and NUGC4-*IQGAP1*^KO^ cells at the levels of transcriptome, mitochondrial proteome and alternative splicing (**Figure S2C**). Interestingly, a single component of the mitochondrial respiratory chain complex I were found to be deregulated at more than one level (mRNA, protein and AS). This was *NDUFS4* (NADH:Ubiquinone Oxidoreductase Subunit S4), an accessory subunit of mitochondrial respiratory chain complex I. According to the RNA seq analysis, *IQGAP1*^KO^ leads to increased skipping of exon 2 in the *NDUFS4* mRNA, predicted to result in ORF disruption and generation of an NMD substrate (**Figure 2A, B**). This event is the most confident hit in our analyses for AS changes based on the number of junction reads (**Figure 2C**). We should stress the point that in NUGC4 cells, exon 2 is 100% included, and skipping is only detected in the absence of IQGAP1. This change in the AS pattern agrees with the gene expression and proteomic analyses, showing down-regulation of *NDUFS4* transcript and protein levels in NUGC4-*IQGAP1*^KO^ cells compared to the parental ones. We validated the results of the RNA seq analysis and confirmed the increase in exon 2 skipping upon IQGAP1 depletion in NUGC4 cells (**Figure 2D**), as well as the downregulation of *NDUFS4 mRNA* expression (**Figure 2E, S2E**). We also confirmed the specific reduction of NDUFS4 protein levels both in whole cell lysates and mitochondrial enriched fractions of NUGC4-*IQGAP1*^KO^ cells compared to the parental ones which agrees with the predicted ORF disruption caused by the altered AS of *NDUFS4* pre-mRNA (**Figure 2F**). This change in exon 2 inclusion was not detected in MKN45 and in HFE145 upon IQGAP1 KO (**Figure S2D**) nor were the downregulation of NDUFS4 at the mRNA or protein levels (**Figure S2E-F**).

**Figure 2.**
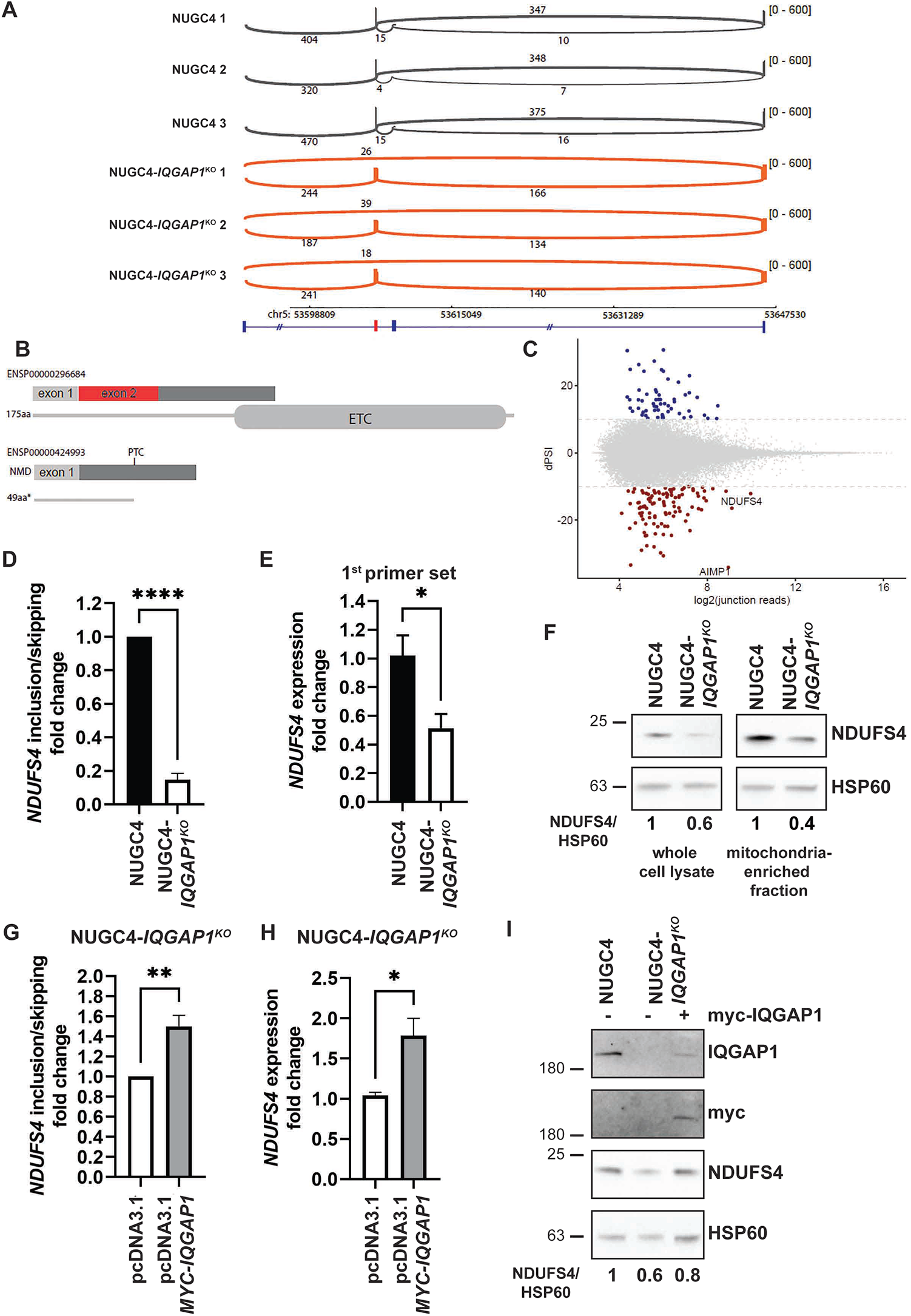
IQGAP1 regulates AS and expression of the mitochondrial respiratory chain complex I subunit NDUFS4. A. Sashimi plot for the visualization of the NDUFS4 AS event derived from aligned RNA seq data from NUGC4 and NUGC4-*IQGAP1*^*KO*^ cells. Exon-exon junction reads are indicated by the numbers above the line. The chromosomal location is indicated below. At the bottom, a schematic shows the exons and the introns involved or surrounding the alternative event. The box in red indicates the alternative exon. B. Schematic showing the predicted impact that the alternative splicing event has on the NDUFS4 protein. ETC, Electron Transport Chain; PTC, Premature Termination Codon. C. The number of reads of the RNA seq data were plotted against the dPsi values for all events between control and KO cells. We highlight the 2 most confident AS changes affecting NDUFS4 and AIMP1. D. RT-qPCR analysis of NDUFS4 exon 2 inclusion versus skipping, using isoform-specific primers. Biological triplicates were used for each cell line. E. RT-qPCR analysis of the same samples as in (D) to monitor NDUFS4 expression, using a set of primers that recognize both the inclusion and exclusion isoforms. GAPDH levels were used for the normalization. F. Western blot analysis of NDUFS4 levels in whole cell and mitochondria-enriched fractions from NUGC4 parental and KO cells. The mitochondrial matrix marker HSP60 was used as loading control. An experiment representative of 4 independent experiments is presented (see also Figure S2A for lysate preparations), where NDUFS4 levels are expressed relatively to the control (NUGC4). G. Same as (D) for NUGC4-*IQGAP1*^KO^ cells transfected with pcDNA3.1 or myc-IQGAP1 expressing plasmid. The mean of the 3 independent experiments is presented. H. Same as (E) for NUGC4-*IQGAP1*^KO^ cells transfected with pcDNA3.1 or myc-IQGAP1 expressing plasmid. The mean of the 3 independent experiments is shown. I. NDUFS4 levels in whole cell lysates of NUGC4 and NUGC4-*IQGAP1*^KO^cells transfected with pcDNA3.1 and NUGC4-IQGAP1KO cells transfected with myc-IQGAP1 expressing plasmid were analyzed by Western blot. HSP60 was used as loading control. Anti-IQGAP1 and anti-myc immunoblotting was used for the detection of endogenous and exogenous IQGAP1. One experiment representative of 3 replicates is presented and NDUFS4 levels are expressed relatively to the control (NUGC4). Error bars represent ±SEM. *, p<0.05; **, p<0.01; ****, p<0.0001. Numbers indicate MW in kDa.

To directly address the connection between IQGAP1 and the *NDUFS4* exon 2 skipping event we transfected NUGC4-*IQGAP1*^*KO*^ cells with a plasmid expressing a Myc-tagged IQGAP1 and assayed the inclusion of exon 2 by RT-qPCR (**Figure 2G**). The splicing pattern of *NDUFS4* exon 2 was partially rescued in IQGAP1 expressing cells. This rescue was also reflected at the mRNA and protein levels of NDUFS4 (**Figure 2H, I**). Taken together these data indicate the involvement of IQGAP1 in the regulation of NDUFS4 levels via control of its AS.

### Altered NDUFS4 levels in IQGAP1^KO^ cells result in deficient mitochondrial complex I assembly and function

NDUFS4 is an accessory subunit of mitochondrial respiratory complex I with an essential role in its assembly and stability. Mitochondrial respiratory complex I is assembled from pre-formed modules (N, Q/P_p_-a, P_p_-b, P_d_-a and P_d_-b) in a step-wise manner, and NDUFS4 is incorporated during the final steps of the process and likely mediates N-to-Q-module attachment^21,22^. NDUFS4 is a hotspot for mutations that cause autosomal recessive Leigh syndrome, a lethal mitochondrial disease that primarily affects the Central Nervous System, and is characterized by complex I deficiency. NDUFS4 deficiency in an *NDUFS4*^KO^ mouse model and Leigh patient fibroblasts is known to result in reduced attachment between the N and Q modules, reduction of the steady state levels of several core subunits of complex I and increased stabilization of a NDUFAF2 (NADH:Ubiquinone Oxidoreductase Complex Assembly Factor 2) -Pp-b +Q-Pp-a intermediate^22^. In our analyses and in agreement to these observations, the protein levels of several core complex I subunits, in particular of the N- and Q-modules, such as NDUFS1 (NADH-ubiquinone oxidoreductase 75 kDa subunit; **Figure 3A, S3A** and **Table S4**) are significantly lower in NUGC4-*IQGAP1*^*KO*^ compared to NUGC4 cells. The levels of subunits of other mitochondrial complexes were not affected by IQGAP1 depletion. To assay mitochondrial complex I assembly in the absence of IQGAP1, we prepared crude mitochondrial fractions and solubilized them with digitonin that preserves the complex I (CI) monomers, but also the super-complexes (SC) with complex III (CIII) and IV (CIV) or solubilized with DDM that maintains only CI monomers^23^. The fractions were analyzed by Blue Native Gel Electrophoresis, followed by immunoblotting against the complex I components NDUFS4, NDUFS1 (**Figure 3B, C**). A lower amount of NDUFS4-containing CI was obviously detected in NUGC4-*IQGAP1*^*KO*^ cells, especially in the CI monomer fractions. In addition, deficient complex I assembly was evident in NUGC4-*IQGAP1*^*KO*^ cells, where a lower band, detected by the antibody against NDUFS1, was present in both DDM and digitonin solubilized mitochondrial fractions, presumably corresponding to the N-module based on its size and the presence of NDUFS1 (**Figures 3B, C and S3B**). In addition, we observed the stabilization of an NDUFAF2-containing intermediate specifically in NUGC4-*IQGAP1*^*KO*^ cells (**Figure S3B**). Such deficient complex I assembly was not detected in *IQGAP1*^KO^ cells where the altered exon 2 skipping of *NDUFS4* has not been detected (**Figure S3B** for HFE145 and HFE145-*IQGAP1*^*KO*^).

**Figure 3.**
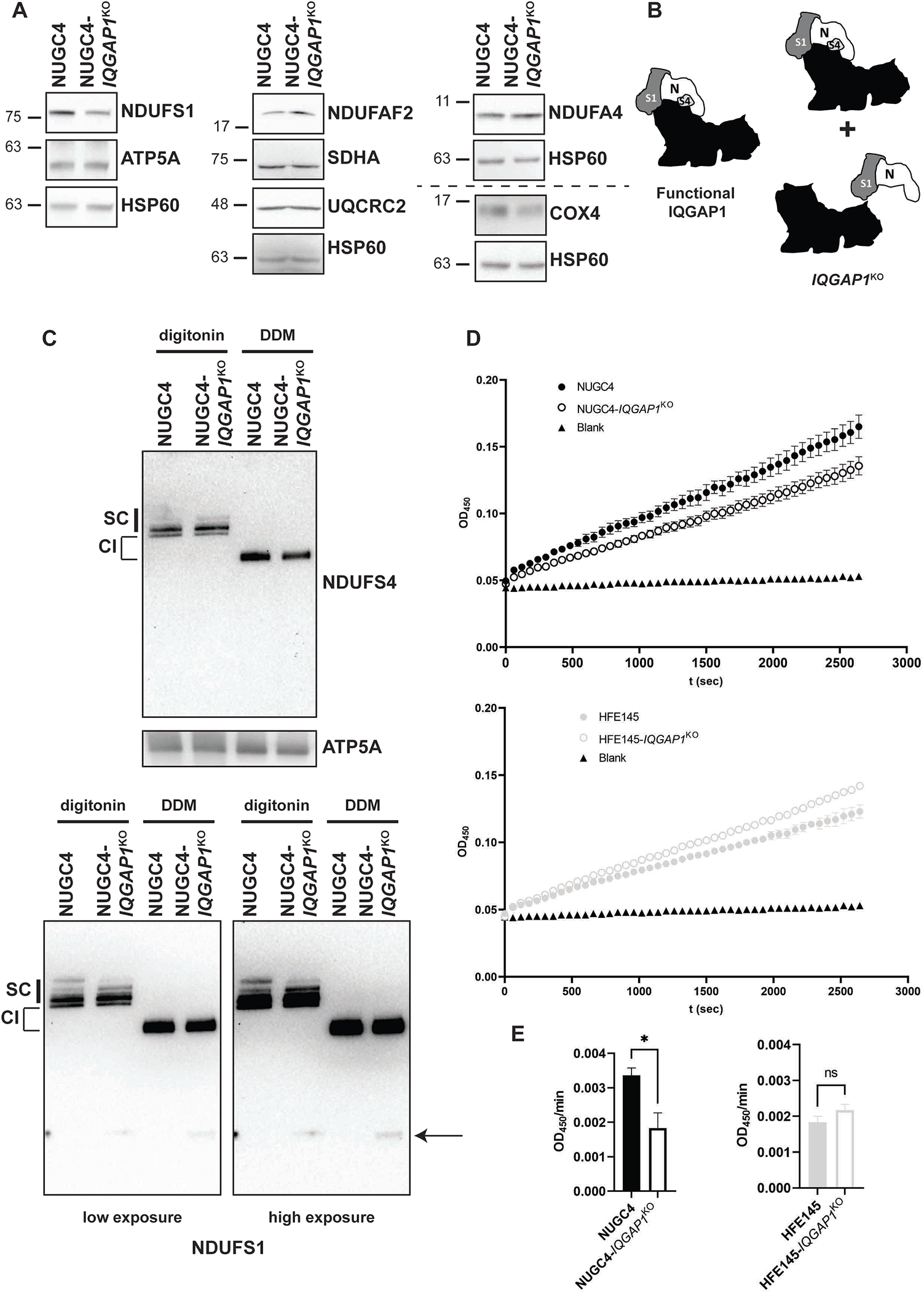
Deregulation of NDUFS4 AS impairs mitochondrial respiratory chain complex I assembly and activity. A. Western blot analysis of mitochondria-enriched fractions from NUGC4 and NUGC4-*IQGAP1*^KO^ cells, using antibodies against different subunits of mitochondrial respiratory chain complexes. Complex I: NDUFS1, NDUFAF2; Complex II: SDHA (Succinate Dehydrogenase Complex Flavoprotein Subunit A); Complex III: UQCRC2 (Ubiquinol-Cytochrome C Reductase Core Protein 2); Complex IV: NDUFA4 (NDUFA4 mitochondrial complex associated), COX4 (Cytochrome c oxidase subunit 4); Complex IV: ATP5A (ATP synthase lipid-binding protein). HSP60 levels were used for normalization. Numbers indicate MW in kDa. B. Schematic representation of Complex I and its intermediates detected in NUGC4 and NUGC4-*IQGAP1*^KO^ cells as presented in (C). N, N-module; S1, NDUFS1; S4, NDUFS4. The Q/P subcomplex of Complex I is represented by the black area. C. BNGE analysis of mitochondrial respiratory chain complexes from NUGC4 and NUGC4-*IQGAP1*^KO^ cells. Crude mitochondria fractions were solubilized in digitonin, which maintains the interaction of complex I (CI) in association with complex III and IV (supercomplexes, SC), or with n-Dodecyl-B-D-Maltoside (DDM), which allows the efficient extraction of CI as monomer. Note that the migration pattern of CI monomers differs depending on the detergent used for the solubilization^38^. The arrow indicates the NDUFS1-containing intermediate which accumulates only in NUGC4-*IQGAP1*^KO^ cells. Complex V levels, detected with an anti-ATP5A antibody, were used to monitor loading. C. Mitochondrial complex I activity expressed as Optical Density (OD) at 450nm was plotted against time for NUGC4 vs NUGC4-*IQGAP1*^KO^ (up) and HFE145 vs HFE145-*IQGAP1*^KO^ (bottom) cells. The curve corresponds to a single experiment representative of 3 (NUGC4) or 2 (HFE145) replicates. D. Mitochondrial complex I activity of NUGC4 vs NUGC4-*IQGAP1*^KO^ (left) and HFE145 vs HFE145-*IQGAP1*^KO^ (right) whole cell lysates, expressed as the mean slope of the curves presented in (C). Error bars represent ±SEM. *, p<0.05.

Mitochondrial complex I deficiency in IQGAP1 KO cells was verified by measuring complex I activity as an electron donor^24^. Using either whole cells or isolated mitochondria, significant reduction in the catalytic activity of Complex I was detected in the NUGC4 cells lacking IQGAP1 compared to the parental cells (**Figure 3C-D and S3C**). Interestingly, in HFE145 cells where IQGAP1 is dispensable for NDUFS4 exon 2 inclusion, we did not detect its necessity for maximal Complex I activity (**Figure 3C-D**). Thus, IQGAP1 KO in NUGC4 cells leads to deficient mitochondrial complex I assembly and function.

### Mitochondrial physiology is affected in gastric cancer cells upon IQGAP1 KO

To our knowledge, there are no reports on a direct involvement of IQGAP1 in mitochondria homeostasis. To address a possible general involvement of IQGAP1 in regulating mitochondrial abundance, we assessed mtDNA content **(Figure S4A**), mitochondrial mass (**Figure S4B**) and expression levels of transcription factors involved in mitochondrial biogenesis (mitochondrial transcription factor A-TFAM, peroxisome proliferator-activated receptor gamma co-activator-PGC1α and nuclear respiratory factor 1-NRF1, **Figure S4C**) in NUGC4 and NUGC4-*IQGAP1*^KO^ cells. No difference was found between parental and KO cells, indicating that IQGAP1 depletion does not impair basal mitochondrial biogenesis and turnover.

Mitochondrial respiratory complex I deficiency has been associated with the deregulation of mitochondrial network morphology and mitochondrial ROS (mtROS) production^25^. We monitored mitochondrial network morphology in *IQGAP1*^KO^ and parental cells by fluorescence microscopy, after immunostaining against the mitochondrial matrix protein HSP60. As presented in **Figure 4A**, IQGAP1 depletion leads to a 30% increase in the percentage of cells with fragmented/aggregated mitochondrial networks at the detriment of cells with tubular or hyperfused networks (**Figure S4D**).

Increased mitochondrial fragmentation is also supported by the measurement of specific parameters, such as Aspect Ratio (AR), reflecting the degree of mitochondrial elongation, and Form Factor (FF), reflecting the degree of mitochondrial network complexity and branching, and the number of identified mitochondria per cell, showing a significant decrease in *IQGAP1*^KO^ cells compared to the parental ones (**Figure 4B-C, S4E**). Importantly, this phenotype was largely rescued when cMyc-IQGAP1 was introduced in the *IQGAP1*^*KO*^ cells (**Figure 4B-C, S4E**). The fragmented mitochondrial network finding was accompanied by a decrease in mtROS production in KO compared to the parental NUGC4 cells (**Figure 4D**), which was also rescued upon expression of Myc-IQGAP1 in the cells (**Figure 4E**), supporting the deficiency in mitochondrial function in the absence of IQGAP1.

**Figure 4.**
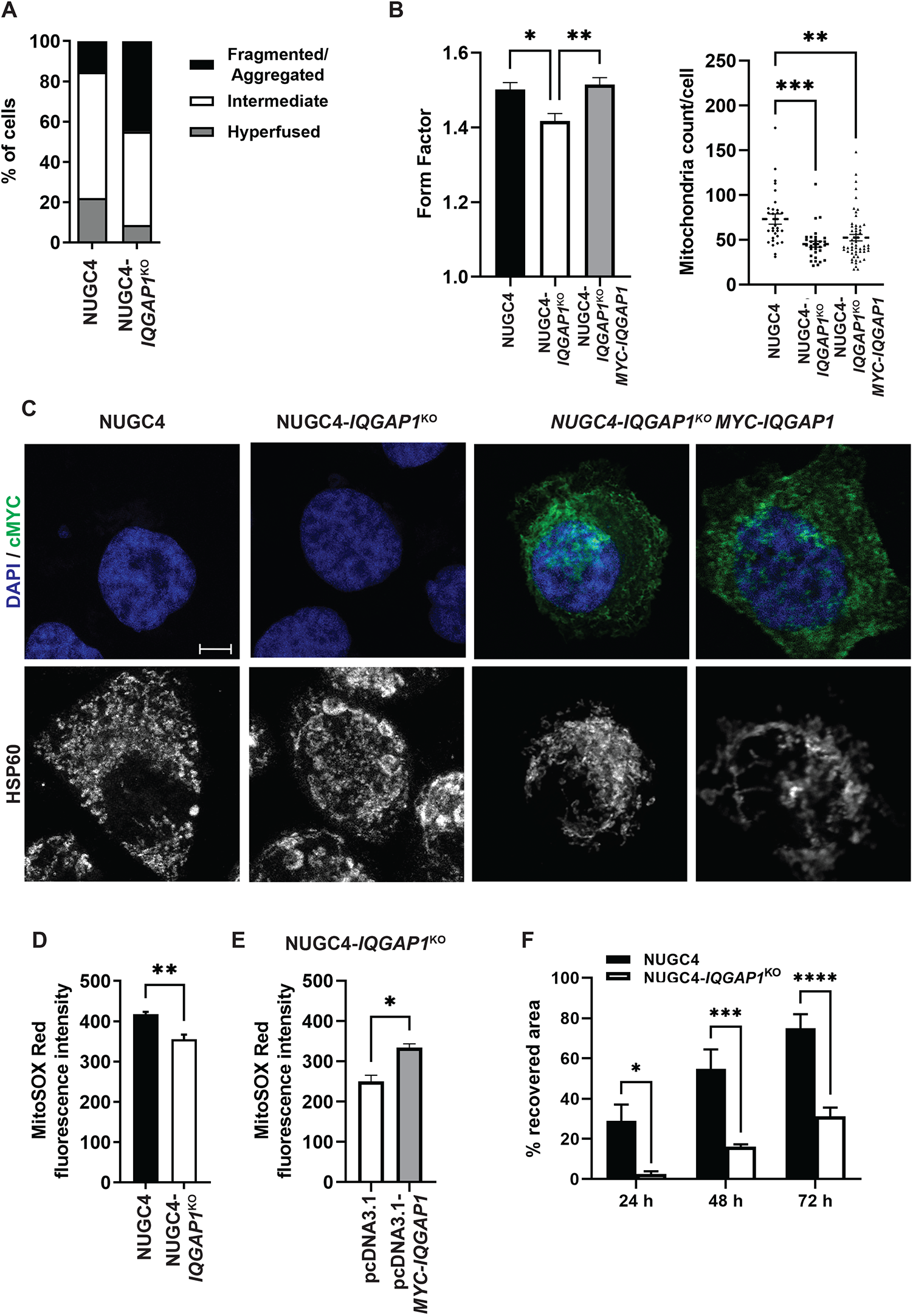
IQGAP1 depletion recapitulates mitochondrial defects and migration impairment of NDUFS4^KO^ cells. A. Relative distribution of the distinct mitochondrial network morphology phenotypes (hyperfused, tubular and fragmented/aggregated, see also Figure S4C) in NUGC4 and NUGC4*-IQGAP1*^*KO*^ cells. Immunostaining against HSP60 was used to visualize mitochondrial networks. B. Form Factor and mitochondria count per cell were measured for NUGC4 and NUGC4-*IQGAP1*^KO^ cells transfected with pcDNA3.1 and NUGC4-*IQGAP1*^KO^ cells transfected with pcDNA3.1-MYC-IQGAP1, immunostained against MYC and HSP60. At least 30 cells per condition were used for the analysis. C. Representative confocal microscopy images of the samples presented in (B). DAPI staining was used to visualize the nuclei. The scale bar is at 5μm. D. MitoSOX Red fluorescence of NUGC4 and NUGC4-*IQGAP1*^*KO*^ cells measured by flow cytometry analysis. The mean fluorescence intensity of triplicate experiments is presented. E. The same as in (D) for NUGC4-*IQGAP1*^KO^ cells transfected with pcDNA3.1 and pcDNA3.1-MYC-IQGAP1. The mean of 2 independent experiments is shown. F. Wound healing assay for NUGC4 and NUGC4-*IQGAP1*^KO^ cells. The migration of the cells was measured 24, 48 and 72 h after wound formation. One experiment representative of 2 is presented. Error bars represent ±SEM. *, p<0.05; **, p<0.01; ***, p<0.001;****, p<0.0001.

It has been reported that NDUFS4 down-regulation impairs the migration of gastric cancer cells^26^. We compared NUGC4 and NUGC4-*IQGAP1*^*KO*^ cells in a wound healing assay and found more than 50% reduction in the migration rate of the KO cells compared to the parental ones (**Figure 4F, S4F**). Of note, IQGAP1 depletion did not impair migration of MKN45 cells (Birladeanu et al. 2021), in which NDUFS4 AS and expression were also unaffected (**Figure S2D-F**). Overall, our data point towards a direct effect of IQGAP1 on the mitochondrial respiratory chain, at least partially through the regulation of AS of NDUFS4, the deregulation of which can be linked to other mitochondrial defects (mitochondrial fragmentation/aggregation, reduction of mitochondrial ROS production) observed in the KO cells.

## Discussion

In the present study we describe a new level of regulation of mitochondrial function, via alternative splicing. The nucleus is known to regulate mitochondrial function, a regulation mainly based on the expression of nuclear-encoded mitochondrial proteins that induce mtDNA gene expression, and on several transcription factors and co-regulators that control the expression of the nuclear-encoded mitochondrial proteome. This anterograde communication can also signal the existence of nuclear stress to mitochondria, thereby bringing about a reduction in mitochondrial metabolism^1^. With our work we add one more layer of anterograde communication between the nucleus and the mitochondria, this of AS. We present evidence that the pre-mRNA of the mitochondrial Complex I component *NDUFS4* can be alternatively spliced in gastric cancer cells and the change of its AS pattern depends on IQGAP1, a scaffold protein known as a cellular integrator of many signalling pathways.

Complex I (CI) is the largest multiprotein enzyme complex of the respiratory chain (∼1 MDa) consisting of three functional modules: the N-module (NADH binding and oxidation), the Q-module (electron transfer to ubiquinone) and the P-module (proton pumping). CI consists of 45 subunits, 37 encoded by the nuclear genome and 7 subunits by the mitochondrial genome (mtDNA). Biogenesis of CI is assisted by at least 14 nDNA encoded assembly factors and its assembly occurs by a mechanism in which five CI protein modules are pre-formed (N, Q/Pp-a, Pp-b, Pd-a and Pd-b) and subsequently combined. Mutations in genes coding for these structural CI subunits or in genes involved in its assembly have been associated to CI deficiency and are primarily associated with Leigh syndrome (LS). The NDUFS4 gene encodes the 18-kDa NDUFS4 accessory subunit of CI, which plays an important role during CI assembly and in CI stability. Various *Ndufs4*^*−/−*^ mouse models have been generated to study the LS mechanism and have shown that the clinical phenotype associated with *Ndufs4* KO is primarily due to brain abnormalities^27^. Studies using either immortalized mouse embryonic fibroblasts^25^ or primary fibroblasts from patients carrying mutations in *NDUFS4* showed that loss of NDUFS4 affects complex I assembly and subsequently activity and causes complex I instability, accumulation of a partially assembled 830 kDa Complex I subcomplex and weak binding of the N-module. In all KO mouse tissues, NDUFS4 loss leads to near complete absence of NDUFA12, increased levels of NDUFAF2, and reduction of many complex I subunits by mitochondrial proteases to maintain mitochondrial integrity and prevent production of ROS. Even though CI deficiency has been connected to accumulation of ROS, studies with immortalized embryonic fibroblasts from KO mice, and fibroblasts from patients carrying a PTC introducing mutation in *NDUFS4* that results in complete loss of the protein showed no ROS accumulation due to the NDUFS4 deficiency. Furthermore, NDUFS4 deficiency has been connected to altered mitochondrial morphology in primary fibroblasts isolated from mouse muscle fibers and skin tissue and patients’ samples^22^.

In IQGAP1 KO gastric cancer cells, we have detected increased skipping of exon 2 of *NDUFS4* pre-mRNA. This results in introduction of a PTC in the mRNA and subsequent downregulation of it and the resulting protein. Exon 2 skipping due to mutations have been described in LS patients and result in reduction of fully assembled CI^28–30^. Our analyses of mitochondrial phenotypes that have been connected to NDUFS4 deficiency were in good agreement with the literature: we detected a partially assembled CI, with reduced binding of the N-module and the reduction of the amount of other CI subunits including NDUFAF12 (**Table S4**). Furthermore, disrupted mitochondrial network and altered morphology, together with reduced complex I activity were apparent in these cells, together with reduced ROS production. This latter finding agrees with studies showing a decreased ROS production occurring in cells harboring CI disassembling mutations^31,32^, even though the literature is still controversial regarding ROS production when CI is completely disassembled. Furthermore, the majority of the existing literature comes from studies on non-cancerous cell types. The molecular mechanisms linking CI to tumour progression and adaptation are still controversial. CI-related mitochondrial dysfunction has been recognized as a major contributing factor in cancer. In particular, xenograft studies have shown that when mtDNA mutations compromise both CI activity and assembly, tumour growth is decreased or even inhibited upon injection of the mutated cells in immunodeficient mice^33,34^. A very recent study pinpointed to NDUFS4 protein as a prognostic marker in gastric cancer. Reduction of NDUFS4 levels by siRNAs resulted in reduced migration and invasion of gastric cancer cells and reduced tumour volume in xenograft models, which agrees with our findings on the IQGAP1 KO cells, where NDUFS4 is alternatively spliced and subsequently the levels of the mRNA and the protein are reduced. Thus, CI-related mitochondrial dysfunction is connected to reduced levels of NDUFS4 and to reduced tumour growth and progression, at least for gastric cancer cells.

Recently, we showed that IQGAP1 is involved in gastric cancer progression via regulation of the AS of a cell cycle regulon^11^. We showed that IQGAP1 interacts with many splicing factors and spliceosome components and it is important for post-translational modifications of at least hnRNPM. In the present work, we report a different role of IQGAP1 in AS regulation. In a gastric cancer cell line that relies largely on mitochondrial respiration, IQGAP1 is involved in mitochondrial function at least by altering the AS of a complex I component. Even though the same interactions between IQGAP1 and splicing factors have been detected in both gastric cancer cell types, IQGAP1 contributes to the regulation of the AS of a completely different subset of AS events in the two cell lines. In a cancer cell line that belongs to the CIN subtype (MKN45)^18^, a subtype characterized by a major deregulation of mitotic pathways, IQGAP1 participates in the control of the cell cycle. In NUGC4, a cell line of the GS subtype that seems to rely much more on mitochondrial respiration, IQGAP1 controls complex I function at least via the AS of NDUFS4. Interestingly, very recently, the role of IQGAP1 in post-translational modifications of splicing factors was reported in HNSCC cells, whereas it seemed that once again, the splicing regulatory role of IQGAP1 is different in the normal human keratinocytes cells tested by Muehlbauer et al^35^. Based on these data, it is possible that IQGAP1 regulates AS in close connection to cancer related signals and possibly not in normal cells. Towards this, attests our observation of a lack of similar phenotypes in a human normal gastric epithelial cell line that lacks IQGAP1 (unpublished observations and data presented in this manuscript). Furthermore, we have previously shown that at least in one gastric cancer cell line, IQGAP1 is important for the connection between heat – induced signals and AS regulation, via participating in the distribution of splicing components in the nucleus, a different way of controlling AS outcome, as it has been previously demonstrated^36,37^. Given the IQGAP1’s known role in a plethora of signalling pathways, we anticipate that its involvement in cancer rather depends on selected signals, which are probably more deterministic in each cancer type. It remains, in a more thorough approach, to detect IQGAP1’s role in splicing regulation not only in gastric cancer or other cancer types, but also in normal cells in relation to specific tumourigenic signals.

## Data Availability

### Funding

InfrafrontierGR/Phenotypos Infrastructure, co-funded by Greece and the European Union (European Regional Development Fund) [NSRF 2014–2020, MIS 5002135]; Hellenic Foundation for Research & Innovation (HFRI) and the General Secretariat for Research and Technology (GSRT) [grant agreement 846 to ZE]; MR was supported by the European Research Council [ERC AdvG 670146]

PmedGR for infrastructure

## Acknowledgements

We thank V. Aidinis, E. Douni, (IFBR, B.S.R.C. “Al. Fleming”) and D.L. Kontoyiannis (Aristotle University of Thessaloniki, Greece) for cell lines and reagents; Sofia Grammenoudi and the Flow cytometry facility of B.S.R.C. “Al. Fleming” for help with transfection experiments and discussions; Vladimir Benes and the EMBL Genecore for RNA-seq analyses; George Stamatakis at the Proteomics Facility of B.S.R.C. “Al. Fleming” for LC-MS/MS analyses and discussions; the personnel of the Imaging facility of B.S.R.C. “Al. Fleming” for help with image acquisition. We also thank Skarlatos G. Dedos (National and Kapodistrian University of Athens, Greece) for reagents, plasmids, discussions and critical reading of the manuscript.

## Notes

### Competing Interest Statement

The authors have declared no competing interest.

### Summary of Updates

The main Figures were missing from the first version, so in this new version this has been corrected

