## Supplementary Figures and Figure and Table legends for "Communication between the nucleus and the mitochondria via NDUFS4 alternative splicing in gastric cancer cells"

**Figure S1.** ***IQGAP1*$^{KO}$ affects differently alternative splicing and metabolic profiles of specific gastric cancer and normal epithelial cell lines. A.** Western blot analysis of IQGAP1 levels in total extract (t.e.), cytoplasmic extract (c.e.), nuclear pellet (n.p.) and nuclear extract (n.e.) from NUGC4 and MKN45 cells. Lamin B1 was used as a marker of nuclear fractions. **B.** Relative distribution of the differentially spliced events between NUGC4 and NUGC4-*IQGAP1*$^{KO}$ cells in the different AS types. Events that are down-regulated in the KO cells are shown in black, and those that are up-regulated are shown in grey. Exon, alternative exon; Alt3, alternative 3' splice site; Alt5, alternative 5' splice site; IR, intron retention. **C.** Respiratory (left) and glycolytic activities of HFE145, NUGC4 and MKN45 cells were measured on a Seahorse analyzer. The mean of 3 independent experiments is presented. **D.** Western blot analysis of IQGAP1 levels in HFE145 and HFE145-*IQGAP1*$^{KO}$ whole cell lysates, using β-actin as loading control. Numbers indicate MW in kDa. **E.** Comparison of respiratory and glycolytic activities of *IQGAP1*$^{KO}$ cells to their parental cells. Each cell line was analyzed in triplicates. Error bars represent ±SEM. *, $p<0.05$; **, $p<0.01$; ****, $p<0.0001$.

**Figure S2.** ***IQGAP1*$^{KO}$ does not perturb *NDUFS4* AS and expression in HFE145 and MKN45 cells. A.** Western blot analysis of total extract (t.e.), cytoplasmic extract (c.e.) and crude mitochondria (c.m.) of NUGC4 and NUGC4-*IQGAP1*$^{KO}$ cells with antibodies against VDAC2 (voltage dependent anion channel 2, outer mitochondrial membrane), HSP60 (heat shock protein 60, mitochondrial matrix) and GAPDH (glyceraldehyde 3-phosphate dehydrogenase, cytosol). Numbers indicate MW in KDa. **B.** Volcano plot of the log2fc change in mitochondrial protein levels between NUGC4 and NUGC4-*IQGAP1*$^{KO}$ cells. Proteins that are significantly down-regulated in the KO cells are marked in blue, whereas those that are significantly up-regulated are marked in red (-log10(p-value)>1.3, log2(fc)>0.8). Significantly down-regulated subunits of mitochondrial respiratory complex I are indicated. **C.** Scatterplot of log2(fc) Gene expression (GE) and log2(fc) Mitochondrial proteome differences between NUGC4 and NUGC4-*IQGAP1*$^{KO}$ cells. In blue are genes that are significantly affected also at the level of AS by *IQGAP1*$^{KO}$. **D.** *NDUFS4* AS pattern was analyzed in parental and *IQGAP1*$^{KO}$ gastric cell lines HFE145 and MKN45 by RT-qPCR using primers specific for the inclusion or exclusion isoform. The mean of biological duplicates is presented. Error bars represent ±SEM. **E.** RT-qPCR analysis of *NDUFS4* expression in *IQGAP1*$^{KO}$ cells and their parental ones, using two different sets of primers that recognize both the inclusion and skipping isoforms. *GAPDH* levels were used for the normalization. The mean of biological triplicates is presented. *, $p<0.05$; ns, not significant. **F.** NDUFS4 levels were analyzed in mitochondria-enriched fractions of HFE145, MKN45 and the derived KO cell lines, using HSP60 as loading control.

**Figure S3. Deregulation of *NDUFS4* AS impairs mitochondrial respiratory chain complex I assembly and activity. A.** Quantitation of respiratory chain complex subunits NDUFS4, NDUFS1, NDUFAF2, SDHA, UQCRC2, COX4, NDUFA4 and ATP5A in mitochondria-enriched fractions of NUGC4 and NUGC4-*IQGAP1*$^{KO}$ cells. HSP60 levels were used for the normalization. The mean of 2-5 independent experiments is presented and the levels of each subunit are expressed relatively to the control NUGC4. In agreement with our proteomic analysis (**Table S4**), NDUFAF2 levels are not down-regulated in KO cells, instead they tend to be higher than in parental cells as previously described for *NDUFS4* null cells[1]. **B.** BNGE analysis of mitochondrial respiratory chain complexes, solubilized with digitonin, from NUGC4, HFE145 and their respective KO cells. Sequential immunoblotting with antibodies against NDUFS1 (CI core subunit), NDUFAF2 (assembly factor, not present in the fully assembled CI) and COX4 (CIV subunit) was performed. Immunoblotting against the complex V subunit ATP5A was used to monitor loading. The arrows indicate the complex I intermediates that accumulate upon $IQGAP1^{KO}$ only in NUGC4 cells. The NDUFS1-containing intermediate migrates below CIV monomers (~200 kDa) and corresponds to free N-module, as confirmed by mass spectrometry analysis (data not shown). The NDUFAF2-containing intermediate migrates just below CI monomers, and most likely corresponds to the 830 kDa intermediate previously described for *NDUFS4*-null mice and patients with Leigh syndrome[1]. SC: supercomplexes of CI with CIII and/or CIV. *: non-specific band. **C.** Mitochondrial complex I activity of NUGC4 *vs* NUGC4-$IQGAP1^{KO}$ mitochondria-enriched fractions. The mean of duplicates for each cell line is presented. Error bars represent ±SEM. **, $p$<0.01; ***, $p$<0.001; ****, $p$<0.0001.

**Figure S4**. **$IQGAP1^{KO}$ does not impair mitochondrial biogenesis**. A. qPCR analysis of mtDNA content in NUGC4 and NUGC4-$IQGAP1^{KO}$ cells. The expression of tRNALeu (mtDNA) were normalized against B2M (nDNA) levels. The mean of biological triplicates is presented. B. MitoTracker Green FM fluorescence of NUGC4 and NUGC4-$IQGAP1^{KO}$ cells measured by flow cytometry analysis. The mean fluorescence intensity of triplicates is presented. MitoTRacker Green FM stains mitochondria regardless of their membrane potential. C. qPCR analysis of the mitochondrial biogenesis transcription factors TFAM, NRF1 and PGC1α in NUGC4 and NUGC4-$IQGAP1^{KO}$ cells. GAPDH levels were used for the normalization. The mean of biological triplicates is presented. Error bars represent ±SEM. D. Representative confocal microscopy images of the different mitochondrial network morphology phenotypes in NUGC4 cells, immunostained against HSP60. E. Representative confocal microscopy images of NUGC4 and NUGC4-$IQGAP1^{KO}$ cells transfected with pcDNA3.1 and NUGC4-$IQGAP1^{KO}$ cells transfected with pcDNA3.1-MYC-IQGAP1, after immunostaining with anti-cMyc and anti-HSP60 antibodies. DAPI staining was used to visualize the nuclei. Scale bar: 5μm. E. Representative images of the wound healing assay presented in Figure 4F for NUGC4 and NUGC4-$IQGAP1^{KO}$ cells at 0 and 72 h after wound formation.

**Table S1**. Alternative splicing changes between NUGC4 and NUGC4-$IQGAP1^{KO}$ cells.

**Table S2.** Proteomic analysis data for the proteins appearing in the heatmap of Figure 1A.

**Table S3.** GO term enrichment analyses as generated by ShinyGo[2] for the proteins of the heatmap in Figure 1A.

**Table S4.** Proteomic analysis data of crude mitochondrial extracts for MKN45, NUGC4 and the respective $IQGAP1^{KO}$ cell lines.

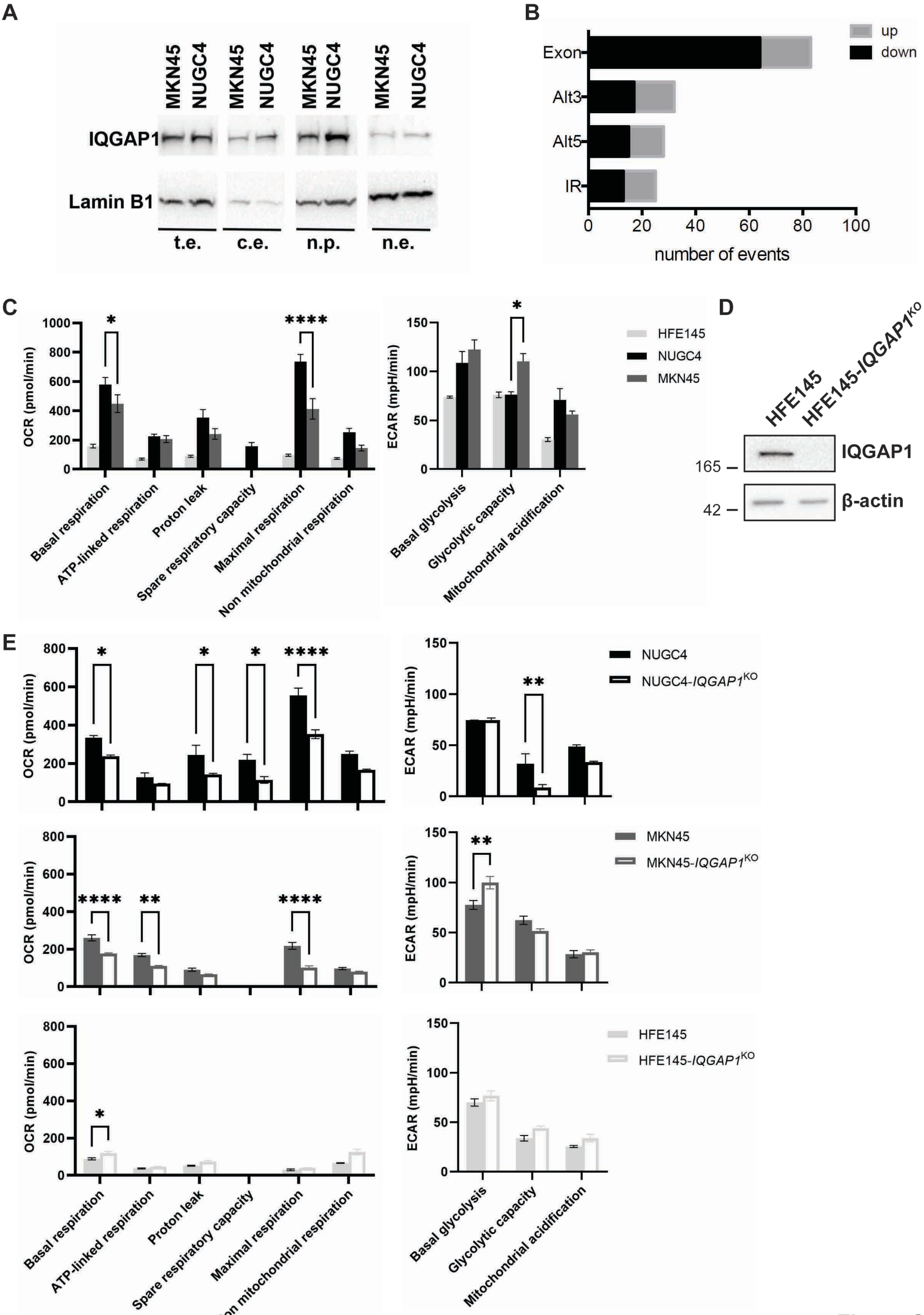
A
MKN45
NUGC4
IQGAP1
Lamin B1
t.e.
c.e.
n.p.
n.e.
B
up
down
Exon
Alt3
Alt5
IR
number of events
C
OCR (pmol/min)
ECAR (mpH/min)
HFE145
NUGC4
MKN45
Basal respiration
ATP-linked respiration
Proton leak
Spare respiratory capacity
Maximal respiration
Non mitochondrial respiration
Basal glycolysis
Glycolytic capacity
Mitochondrial acidification
D
HFE145
HFE145-IQGAP1KO
IQGAP1
β-actin
165
42
E
NUGC4
NUGC4-IQGAP1KO
MKN45
MKN45-IQGAP1KO
HFE145
HFE145-IQGAP1KO

Figure S1

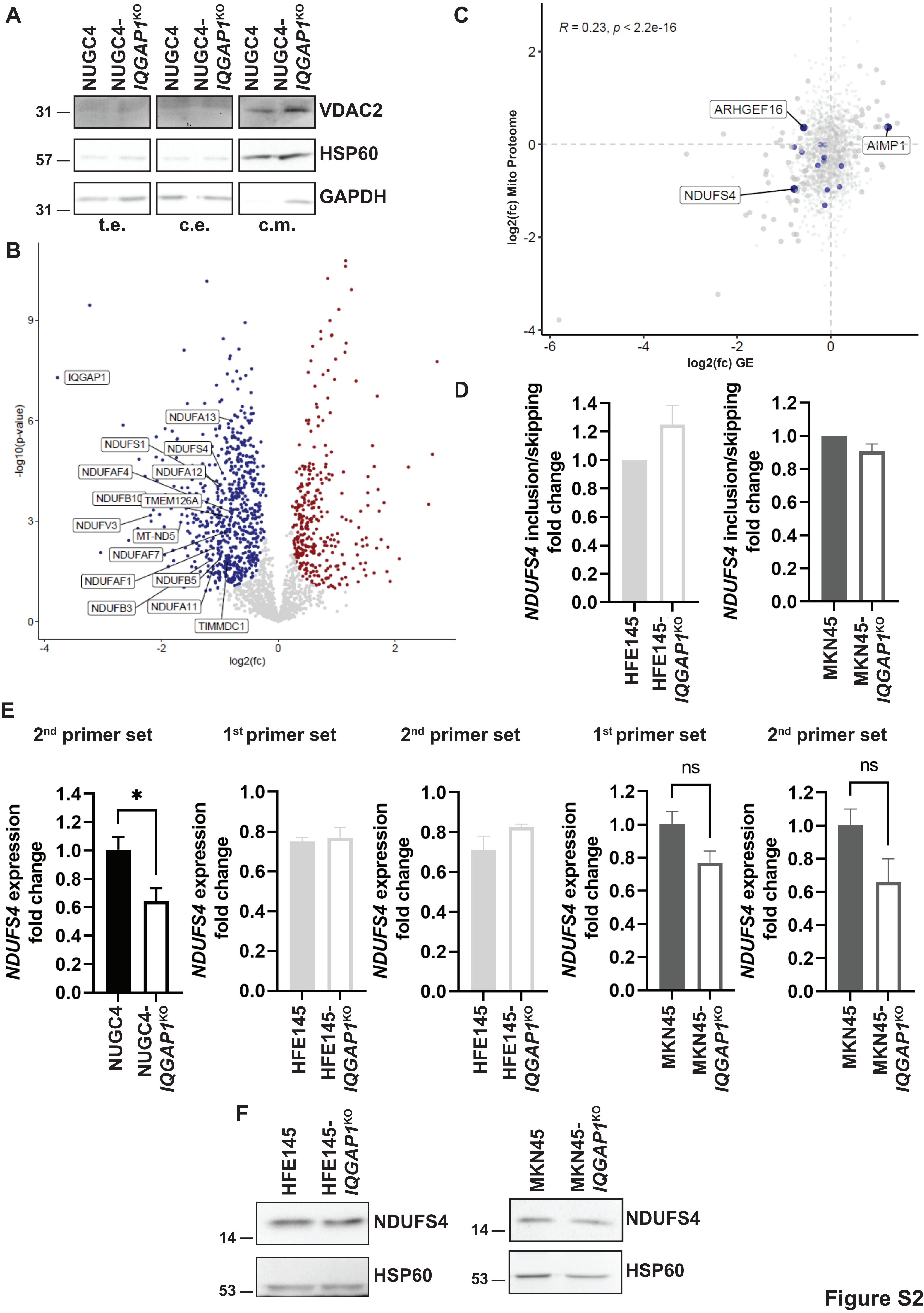
A
NUGC4
NUGC4-IQGAP1KO
31
57
31
VDAC2
HSP60
GAPDH
t.e.
c.e.
c.m.
B
IQGAP1
NDUFA13
NDUFS1
NDUFS4
NDUFAF4
NDUFA12
NDUFB10
TMEM126A
NDUFV3
MT-ND5
NDUFAF7
NDUFAF1
NDUFB5
NDUFB3
NDUFA11
TIMMDC1
-log10(p-value)
log2(fc)
C
R = 0.23, p < 2.2e-16
ARHGEF16
AIMP1
NDUFS4
log2(fc) Mito Proteome
log2(fc) GE
D
NDUFS4 inclusion/skipping fold change
HFE145
HFE145-IQGAP1KO
MKN45
MKN45-IQGAP1KO
E
2nd primer set
1st primer set
2nd primer set
1st primer set
2nd primer set
NDUFS4 expression fold change
*
ns
ns
NUGC4
NUGC4-IQGAP1KO
HFE145
HFE145-IQGAP1KO
MKN45
MKN45-IQGAP1KO
F
HFE145
HFE145-IQGAP1KO
MKN45
MKN45-IQGAP1KO
NDUFS4
HSP60
14
53

Figure S2

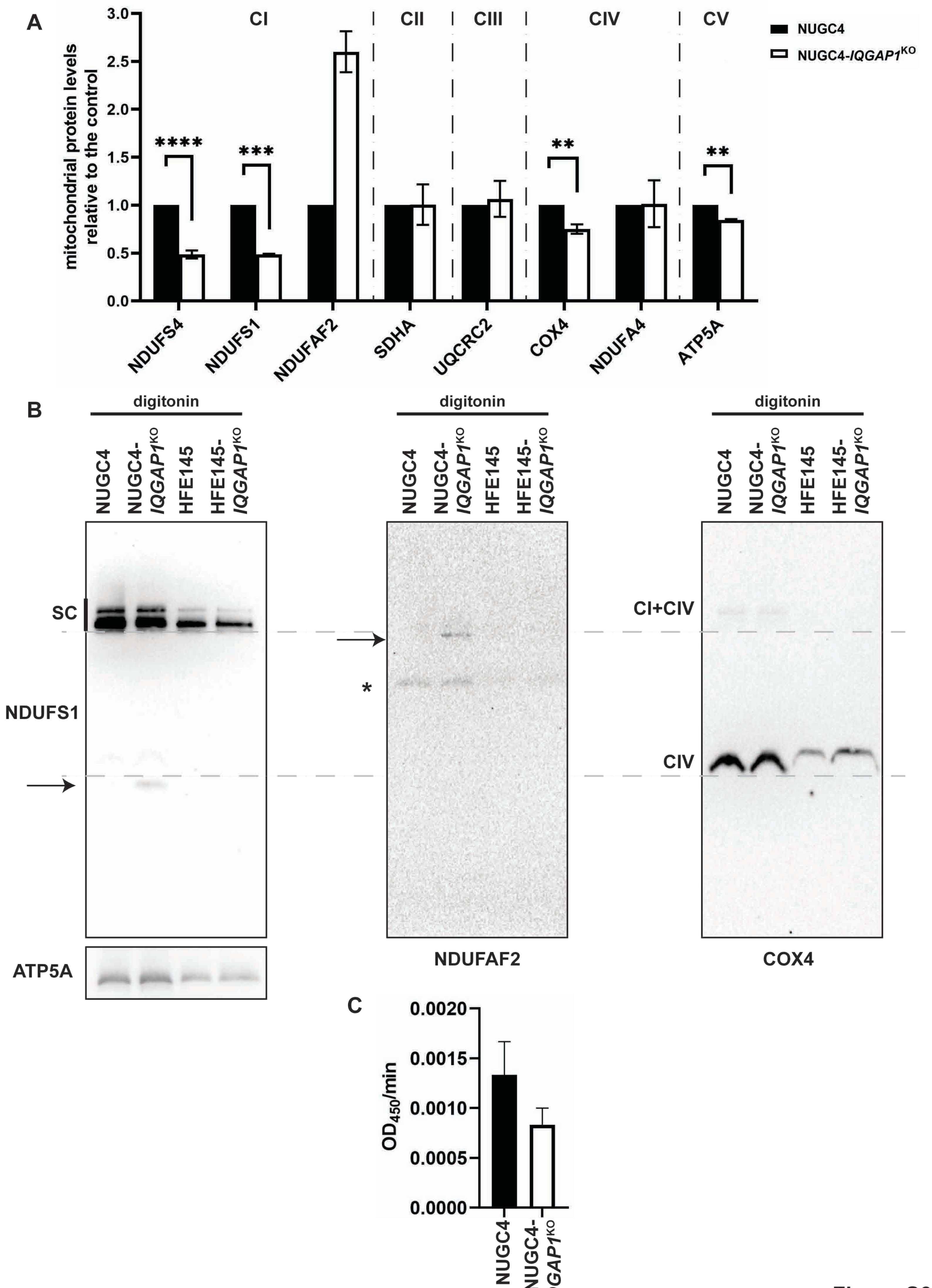
A
B
C
CI
CII
CIII
CIV
CV
NUGC4
NUGC4-IQGAP1KO
mitochondrial protein levels relative to the control
****
***
**
**
NDUFS4
NDUFS1
NDUFAF2
SDHA
UQCRC2
COX4
NDUFA4
ATP5A
digitonin
NUGC4
NUGC4-IQGAP1KO
HFE145
HFE145-IQGAP1KO
SC
NDUFS1
ATP5A
*
NDUFAF2
CI+CIV
CIV
COX4
OD450/min
0.0020
0.0015
0.0010
0.0005
0.0000

Figure S3

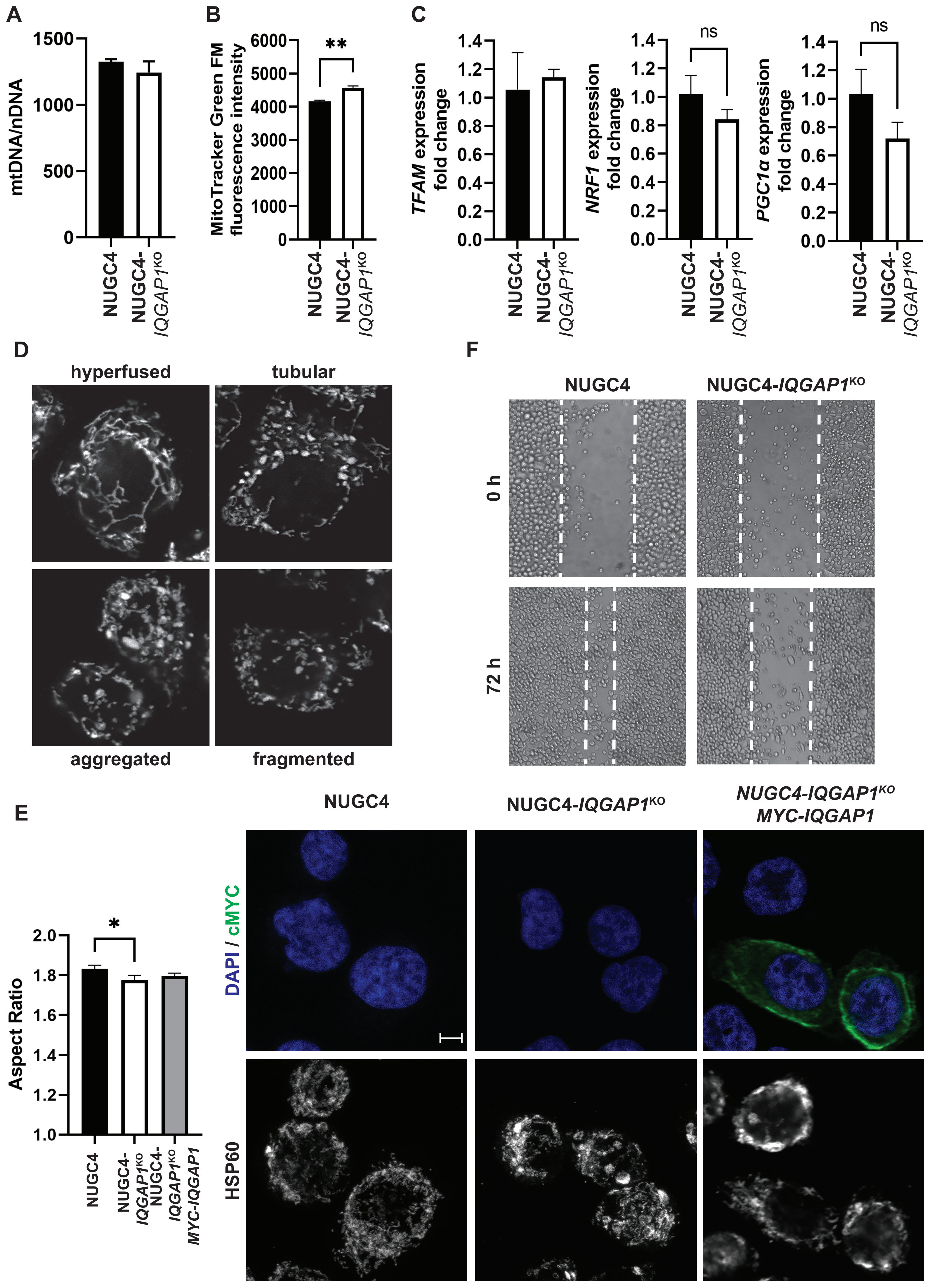
A
mtDNA/nDNA
NUGC4
NUGC4-IQGAP1KO
B
MitoTracker Green FM fluorescence intensity
**
C
TFAM expression fold change
NRF1 expression fold change
PGC1α expression fold change
ns
D
hyperfused
tubular
aggregated
fragmented
F
0 h
72 h
E
Aspect Ratio
*
NUGC4-IQGAP1KO MYC-IQGAP1
DAPI / cMYC
HSP60

Figure S4
