## Supplementary material for "Communication between the nucleus and the mitochondria via NDUFS4 alternative splicing in gastric cancer cells": Table S4

*Table S5. Primers used in this study.*

| Gene | Primer Name | Primer Sequence | Experiment, Figure(s) |
|---|---|---|---|
| *NDUFS4* | NDUF_Incl_F | CCAGGTCGTTGAGGACTTCC | AS analysis, Fig. 2D, 2G, S2C |
| | NDUF_Incl_R | GCGAGCAGGAACAAAGATCC | |
| | NDUF_Skip_F | AGACGTTGTGGCGGAGAAG | |
| | NDUF_Skip_R | AAAGTAGTGATATCCTGGTCGG | |
| | NDUFS4 qPCR_F1 | GTGGTACTGAGGCAGACGTTGT | GE analysis, Fig. 2E, 2F, S2D |
| | NDUFS4 qPCR_R1 | GTCCTGTGCCAATCTCCATGTG | |
| | NDUFS4 qPCR_F2 | TGCTCGCAATAACATGCAGTC | |
| | NDUFS4 qPCR_R2 | GATCAGCCGTTGATGCCCAA | |
| *GAPDH* | GAPDH-F | ACATCAAGAAGGTGGTGAAGCAGG | Normalization GE, Fig. 2E, 2F, S2D, S4B |
| | GAPDH-R | TGTCGCTGTTGAAGTCAGAGGAGA | |
| *PGC1α* | PGC1α-F | CAAGCCAAACCAACAACTTTATCTCT | GE analysis, Fig. S4B |
| | PGC1a-R | CACACTTAAGGTGCGTTCAATAGTC | |
| *NRF1* | NRF1-F | AGGAACACGGAGTGACCCAA | GE analysis, Fig. S4B |
| | NRF1-R | TATGCTCGGTGTAAGTAGCCA | |
| *TFAM* | TFAM-F | CCGAGGTGGTTTTCATCTGT | GE analysis, Fig. S4B |
| | TFAM-R | ACGCTGGGCAATTCTTCTAA | |
| *$tRNA^{LEU}$* | mt-tRNAleu-F | CACCCAAGAACAGGGTTTGT | mtDNA analysis, Fig. 4A |
| | mt-tRNAleu-R | TGGCCATGGGTATGTTGTTA | |
| *B2M* | B2M-F | TGCTGTCTCCATGTTTGATGTATCT | Normalization mtDNA, Fig. 4A, |
| | B2M-R | TCTCTGCTCCCCACCTCTAAGT | |
